# Global computational alignment of tumor and cell line transcriptional profiles

**DOI:** 10.1101/2020.03.25.008342

**Authors:** Allison Warren, Andrew Jones, Tsukasa Shibue, William C. Hahn, Jesse S. Boehm, Francisca Vazquez, Aviad Tsherniak, James M. McFarland

## Abstract

Cell lines are key tools for preclinical cancer research, but it remains unclear how well they represent patient tumor samples. Identifying cell line models that best represent the features of particular tumor samples, as well as tumor types that lack *in vitro* model representation, remain important challenges. Gene expression has been shown to provide rich information that can be used to identify tumor subtypes, as well as predict the genetic dependencies and chemical vulnerabilities of cell lines. However, direct comparisons of tumor and cell line transcriptional profiles are complicated by systematic differences, such as the presence of immune and stromal cells in tumor samples and differences in the cancer-type composition of cell line and tumor expression datasets. To address these challenges, we developed an unsupervised alignment method (Celligner) and applied it to integrate several large-scale cell line and tumor RNA-Seq datasets. While our method aligns the majority of cell lines with tumor samples of the same cancer type, it also reveals large differences in tumor/cell line similarity across disease types. Furthermore, Celligner identifies a distinct group of several hundred cell lines from diverse lineages that present a more mesenchymal and undifferentiated transcriptional state and which exhibit distinct chemical and genetic dependencies. This method could thus be used to guide the selection of cell lines that more closely resemble patient tumors and improve the clinical translation of insights gained from cell line models.

## Introduction

Tumor-derived cell line models have been a cornerstone of cancer research for decades. The genomic and molecular features of over a thousand cancer cell line models have now been deeply characterized ^1^, and recent efforts are systematically mapping their genetic ^2–4^ and chemical ^5^ vulnerabilities. These datasets are thus providing new opportunities to identify potential therapeutic targets and connect these vulnerabilities with measurable biomarkers that can be used to develop precision cancer approaches ^2,5^.

The clinical applicability of results derived from cancer cell lines remains an important question, however, due in large part to uncertainty as to how well they represent the biological characteristics and drug responses of patient tumors. Historically-derived cell line models likely represent an incomplete sampling of the spectrum of human cancers ^6,7^. Many existing models have been propagated for decades *in vitro*, with factors such as clonal selection, cell culture conditions, and ongoing genomic instability all potentially contributing to systematic differences between cell line models and tumors ^7–9^. Furthermore, many cell line models lack detailed clinical annotations. It is therefore critically important to better understand the systematic differences between cell lines and tumors to identify which tumor types have existing cell lines that sufficiently recapitulate their biology and which tumor types do not. Such systematic comparisons may ultimately also help reveal whether patient-derived xenografts, genetically engineered mouse models, or organoid cultures are more, less, or equivalently faithful to human tumors than historical cell lines.

Large datasets such as The Cancer Genome Atlas (TCGA) ^10^ and the Cancer Cell Line Encyclopedia (CCLE) ^1^ include the multi-omic features of approximately 10,000 primary tumor biopsy samples and more than 1,000 cancer cell lines. While the TCGA focuses exclusively on primary tumora (as opposed to metastatic or drug-resistant tumors from which certain cell lines may have been derived) it nevertheless provides a powerful opportunity to begin to perform detailed comparisons of tumors and cell lines systematically across many cancer types. By contrast, previous studies have largely focused on comparing tumors and cell lines within particular cancer types ^11–14^.

In principle, a global alignment of the datasets would allow for the identification of the best cell line models for a given cancer type, without relying on annotated disease labels. Existing global analyses have largely compared samples based on their mutation and copy number profiles ^15^, which are complicated by several factors: a lack of paired normal samples for calling mutations in cell lines, systematic differences in the overall rates of copy number variation and mutations ^13,16–19^, as well as being limited to known cancer-related lesions.

Comparisons based on information-rich gene expression profiles are a promising alternative ^20^, given their demonstrated utility for resolving clinically relevant tumor (sub)types ^21–25^, as well as predicting genetic ^2^ and chemical vulnerabilities of cancer cells ^5,26^. However, a key challenge is that gene expression measurements from bulk tumor biopsy samples are confounded by the presence of stromal and immune cell populations not found in cell lines, often comprising a substantial fraction of the cellular makeup of each sample ^27,28^. Existing approaches for removing the effects of contaminating cells generally require detailed prior knowledge of the expression profiles of each contaminating cell type ^29^, and would not account for other systematic differences between *in vitro* and tumor expression profiles. Furthermore, more general batch effect correction methods typically require either pre-existing subtype annotations, or assume the cell line and tumor datasets have the same subtype composition ^30^.

To address these challenges, we developed Celligner, a method to perform an unsupervised global alignment of large-scale tumor and cell line gene expression datasets. Celligner leverages computational methods recently developed for batch correction of single-cell RNA-seq data and differential comparisons of high-dimensional data, in order to identify and remove the systematic differences between tumors and cell lines, allowing for direct comparisons of their transcriptional profiles. Notably, Celligner aligns pan-cancer gene expression datasets without the need for any additional information (such as tumor type labels, predefined contaminating cell signatures, or tumor purity estimates).

We applied our method to tumor data from TCGA, TARGET, and Treehouse ^31^ and cancer cell line data from CCLE and the Cancer Dependency Map ^1^. This comparison identified cell lines that match well to different tumor subtypes, as well as cancer cell lines that are transcriptionally distinct from their annotated primary cancer types.

## Results

### Alignment of tumor and cell line transcriptional profiles

To illustrate the analytical challenges involved with directly comparing cell line and tumor expression profiles, we first combined several large RNA-Seq gene expression datasets and performed a joint dimensionality reduction analysis. Specifically, we analyzed transcriptional data from 1,249 CCLE cell lines ^1^, 9,806 TCGA tumor samples, 784 pediatric tumor samples from TARGET, and 1,646 pediatric tumor samples from Treehouse ^31^. Although a consistent computational pipeline was used to process all datasets, this analysis revealed a clear separation of cell line and tumor samples (**Fig. 1a**), as expected. This separation was not addressed by applying simple normalization or batch correction methods such as ComBat ^30,32^ (**Supplementary Fig. 1**). This global separation of cell line and tumor samples precludes more detailed assessments of the similarity of samples of different types.

**Figure 1.**
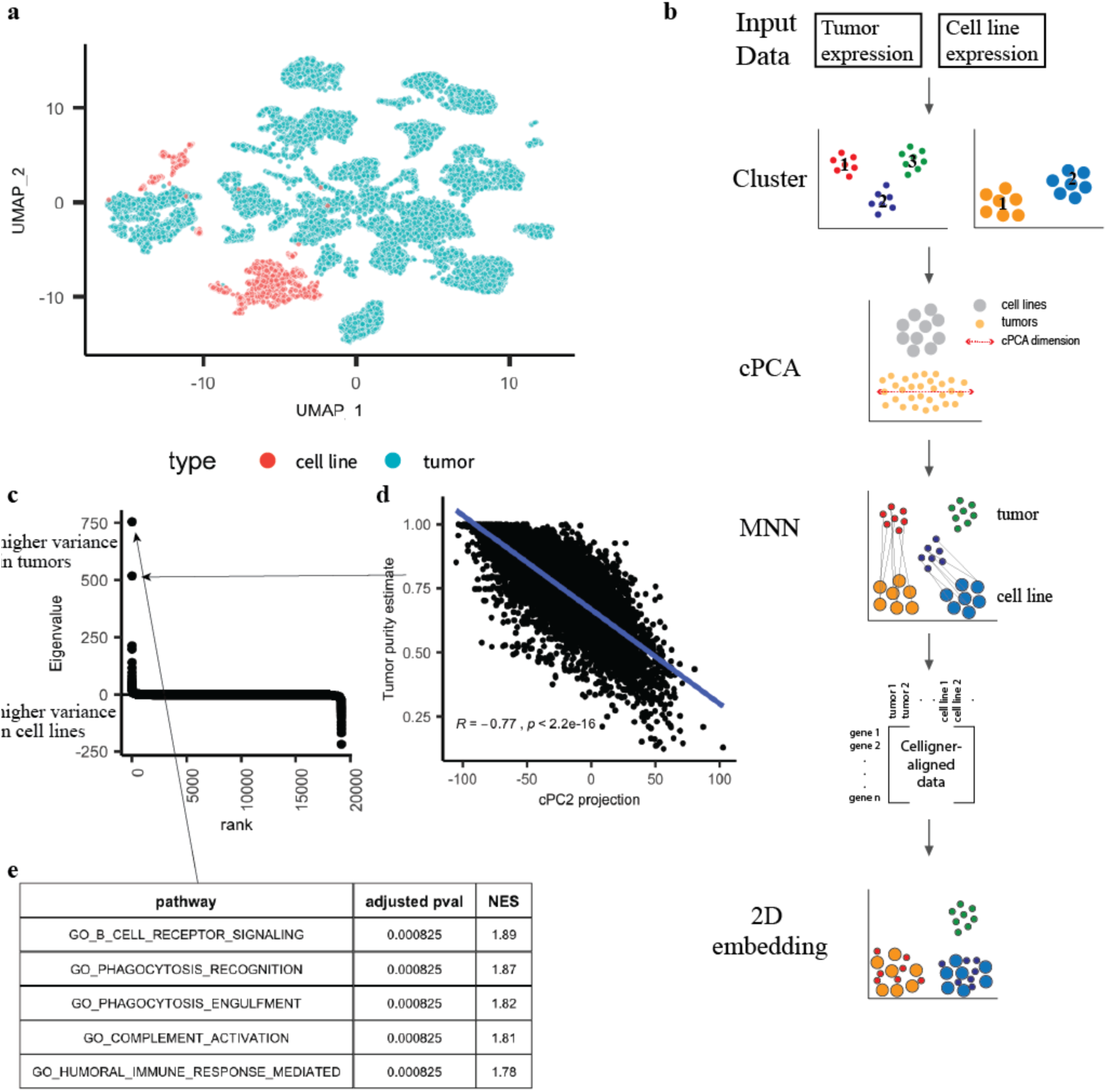
Overview of the Celligner alignment method. **a** A 2D projection of combined, uncorrected cell line and tumor expression data using UMAP. **b** Method: Celligner takes cell line and tumor gene expression data as input, and first identifies and removes expression signatures with excess ‘intra-cluster’ variance in the tumor compared to cell line data using contrastive Principal Component Analysis (cPCA). Then Celligner identifies and aligns similar tumor-cell line pairs to produce corrected gene expression data, using mutual nearest neighbors (MNN) batch correction, which allows for improved comparison of tumors and cell lines. **c** cPC eigenvalues ordered by rank. **d** Correlation between the projection of tumor samples onto cPC2 and their estimated purity (using a consensus measurement of tumor purity). **e** The top five pathways from gene set enrichment analysis (GSEA) of cPC1.

Several features of the problem make alignment of cell line and tumor expression profiles challenging. First, the degree of tumor purity is highly variable across tumor samples ^27,28^, and the transcriptional effects of contaminating normal cells are not captured by a single signature shared across tumor samples and types ^33^. Secondly, differences in the disease (sub)type composition of the datasets can greatly confound attempts at globally aligning the distributions of tumor and cell line expression profiles. Finally, even without the confounding effects of normal-cell contamination, differences between *in vivo* and *in vitro* conditions, as well as technical artifacts, likely give rise to systematic differences in the cancer cells’ gene expression profiles ^34^.

To address these challenges, we developed a multi-step alignment procedure (**Fig. 1b; Methods**). First, to identify gene expression signatures characterizing recurrent patterns of normal cell contamination in tumor samples, we used contrastive principal component analysis (cPCA), a generalization of PCA that uncovers patterns of correlated variation that are enriched in one dataset relative to another ^35^.

Importantly, to avoid biases resulting from the differential disease composition between the two datasets, we first performed an unsupervised clustering of each dataset (**Supplementary Fig. 1; Methods; Supplementary Table 1**), and used cPCA to contrast the intra-cluster covariance structure between the cell line and tumor data. This analysis identified several gene expression signatures with greatly elevated variance across the tumor samples compared to the cell lines (**Fig. 1c**). Gene set enrichment analysis (GSEA) ^36^ of these tumor-specific signatures revealed clear enrichment for immune pathways (**Fig. 1e; Supplementary Fig. 2**), suggesting that cPCA identifies the presence of different contaminating immune cell populations. Furthermore, expression of the second tumor-specific cPC was significantly correlated (*R* = −0.77, *p*-value < 2.2e-16) to independent estimates of tumor purity based on a consensus measurement of tumor purity ^28^, illustrating that this analysis is able to identify multiple independent signatures of contaminating cells (**Fig. 1d**). As the first stage of alignment, we thus removed the top four tumor-specific signatures from both datasets (**Methods**).

While cPCA removes a dominant source of systematic tumor/cell line differences, on its own it does not fully ‘align’ the datasets (**Supplementary Fig. 3**) as it does not account for uniform differences between tumor and cell line profiles of a given disease (sub)type. As a second stage of the alignment, we utilized mutual nearest neighbors (MNN) batch effect correction algorithm to remove the remaining systematic differences between the datasets. MNN batch correction was developed to remove batch effects in single-cell RNA-Seq data. It functions by identifying pairs of samples between datasets where each sample is contained in each other’s set of nearest neighbors in the other dataset, and leverages these MNN pairs to learn a flexible but robust nonlinear alignment of the datasets. Critically, MNN is robust to differences in subtype composition between the datasets, assuming only that the datasets contain a subset of corresponding samples ^37^.

Application of MNN to the tumor and cell line expression profiles identified a set of ‘correction vectors’ (differences in expression profiles between matched tumor/cell line pairs), which on average showed increased expression of immune-related genes, and decreased expression of cell cycle genes, in tumors compared to MNN-matched cell lines (**Supplementary Fig. 3**). While these tumor/cell line differences were largely consistent across samples, the correction vectors also showed patterns that varied across disease types (**Supplementary Fig. 3**), highlighting the importance of using a flexible nonlinear correction method such as MNN to remove such systematic differences. Notably, while MNN on its own provided a broadly similar alignment of the datasets (**Supplementary Fig. 3**), application of cPCA prior to MNN increased the number of MNN pairs identified, and helped mitigate bias towards matching cell lines with higher-purity tumor samples in MNN pairs (**Supplementary Fig. 3**).

We applied this two-stage alignment method (which we refer to as Celligner) to produce an integrated dataset of cell line and tumor gene expression profiles that have been corrected for multiple sources of systematic dataset-specific differences. Indeed, creating a 2D Uniform Manifold Approximation and Projection (UMAP) ^38^ plot with the Celligner-aligned dataset revealed a map of cancer transcriptional profiles with cell line and tumor samples largely intermixed, while still preserving clear differences across known tumor types (**Fig. 2**).

**Figure 2.**
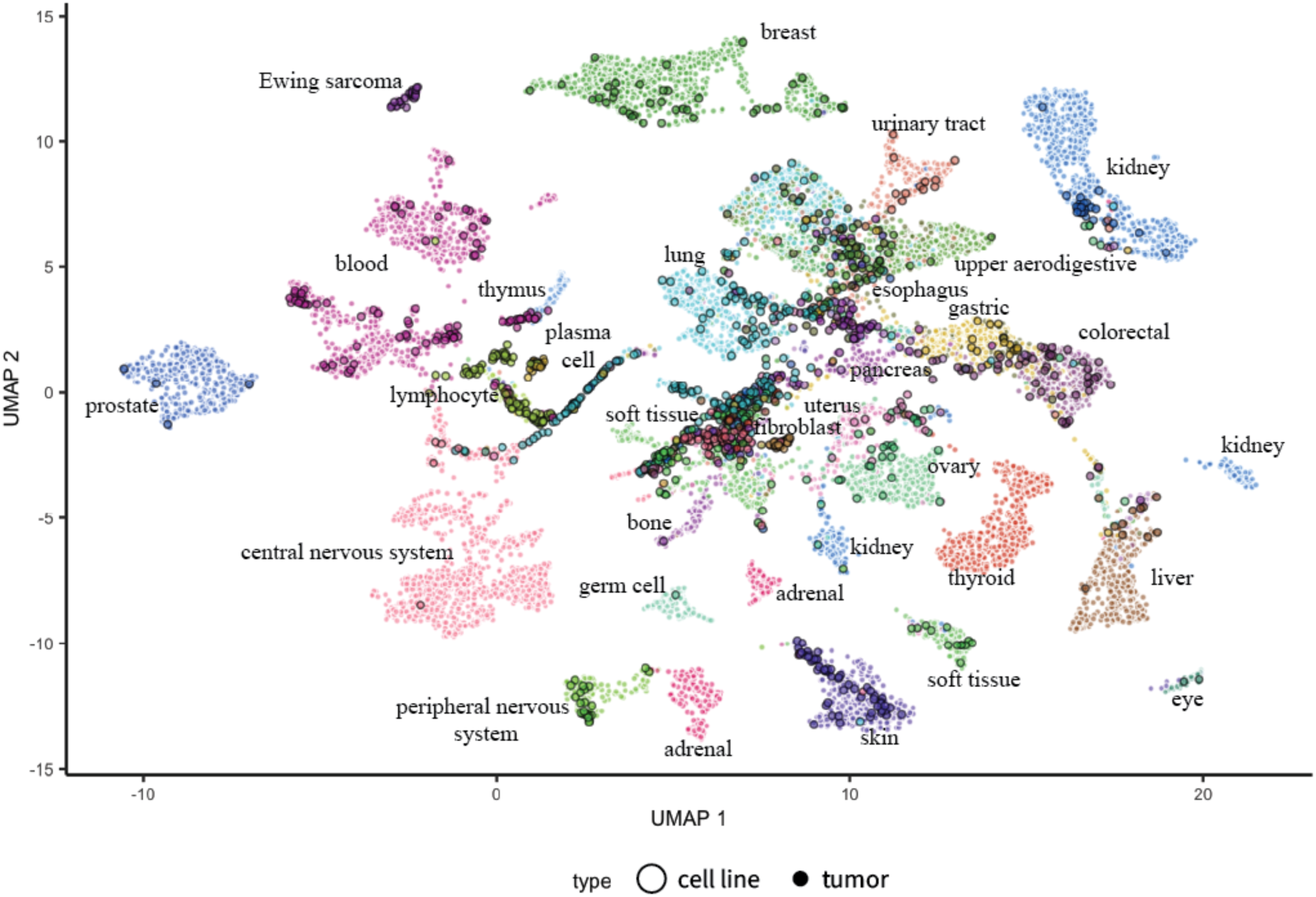
Celligner alignment of tumor and cell line samples. UMAP 2D projection of Celligner-aligned tumor and cell line expression data colored by annotated cancer lineage. The alignment includes 12,236 tumor samples and 1,249 cell lines, across 37 cancer types.

### Alignment preserves meaningful subtype relationships

To evaluate Celligner, we first tested whether it produced an alignment of known disease types and subtypes present in both the tumor and cell line data. As apparent in Figure 2, Celligner removes much of the systematic differences between tumor and cell line expression profiles, producing an integrated map of cancer expression space with clear clusters composed of both cell line and tumor samples. Even though Celligner is completely unsupervised (i.e. does not rely on any sample annotations such as disease type), the aligned tumor and cell line expression profiles largely clustered together by disease type. We quantified this by classifying the most similar tumor type for each cell line, based on its nearest neighbors among the tumor samples. We found that, for disease types found in both datasets, these inferred tumor types matched the annotated cell line disease type 54% of the time (**Fig. 3a; Methods**), while in the uncorrected data the inferred tumor types matched the annotated cell line disease type 45% of the time (**Supplementary Fig. 4**). Celligner correction also increased the measured similarity of tumors and cell lines expression profiles of the same type (**Fig. 3b; Supplementary Fig. 4**).

**Figure 3.**
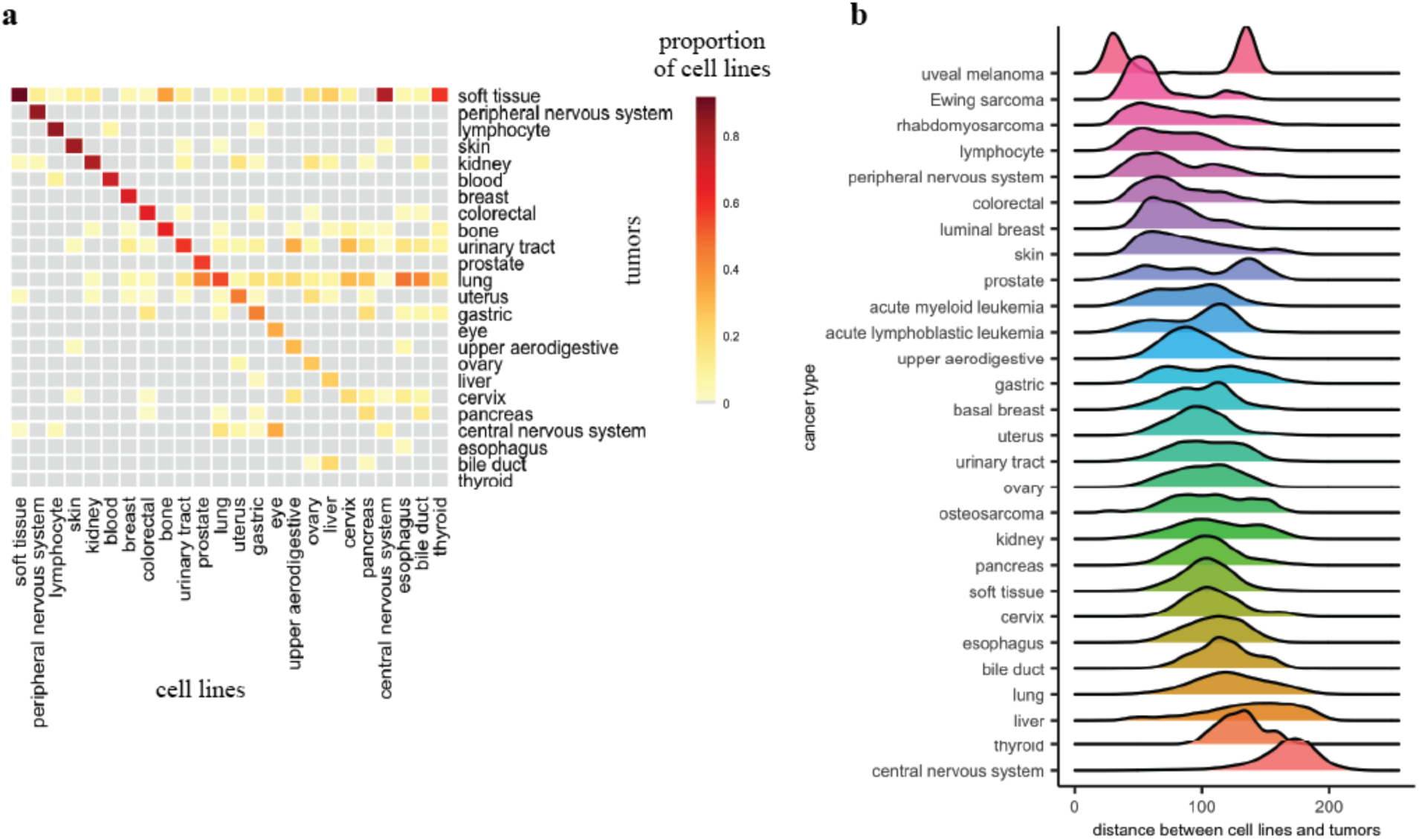
Classification of cell lines by tumor type. **a** The proportion of cell lines that are classified as each tumor type using Celligner-aligned data. **b** Distribution of distances (using top 70 PCs) between cell lines and tumors of the same (sub)type after Celligner-alignment.

A key advantage of Celligner is that it does not assume that all cell line samples in a dataset are necessarily similar to any tumor samples, and vice versa. As a result, we can utilize the Celligner-aligned expression data to identify which cancer types show good agreement between cell lines and primary tumors, and which do not. Although a high proportion of cell lines clustered with tumors of the same cancer type, not all cell lines aligned well with tumor samples. For example, while many soft tissue, skin cancer, and breast cancer cell lines were similar to corresponding tumor samples, we found that central nervous system (CNS) and thyroid cell lines consistently aligned poorly with tumor samples (**Fig. 3a, b**). This observation agrees with previous reports in the literature that *in vitro* media conditions can alter the phenotype of CNS cell lines and cause genomic changes that were not present in the original tumor ^39,40^.

Nevertheless, these analyses illustrate that overall, Celligner tends to group cell lines and tumors of the same disease type. We next sought to determine whether the aligned data also reveal meaningful relationships between more granular subtypes. To this end we aggregated existing subtype annotations for cell line and tumor datasets (**Supplementary Table 1**) ^1,10,41–43^, and found that Celligner also tended to align tumor and cell line samples of the same subtype (**Fig. 4a**). For example, breast cancer tumors and cell lines clustered together by luminal and basal subtypes (**Fig. 4b**). Similarly, the leukemia samples formed clusters that correspond to existing annotations for acute myeloid leukemia (AML) and acute lymphoblastic leukemia (ALL), and the majority of leukemia cell lines aligned to tumors of the same subtype (**Fig. 4c, d**).

**Figure 4.**
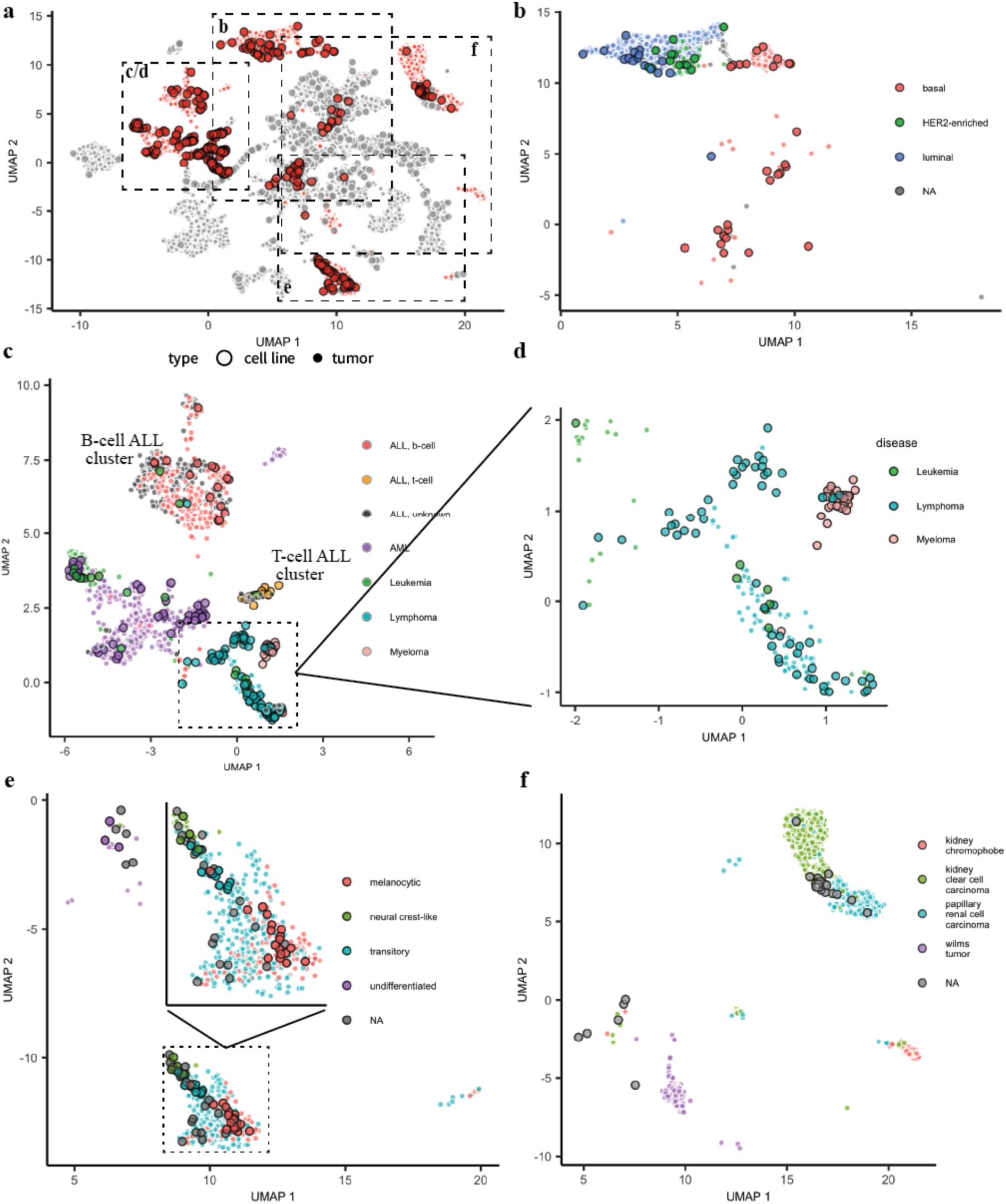
Subtype alignment. **a** UMAP 2D projection of the Celligner-alignment with breast, kidney, hematopoietic, and skin samples highlighted. **b** Luminal and basal breast cancer subtypes cluster together for cell lines and tumors. **c** Leukemia tumor and cell lines subtypes co-cluster in the aligned data. **d** Plasma cell (multiple myeloma) cell lines cluster near the hematopoietic samples, but distinct from the tumors. **e** Melanoma melanocytic, transitory, neural crest-like, and undifferentiated subtypes align between cell lines and tumors. **f** Kidney chromophobe tumor samples do not cluster with any cell lines.

We also tested whether Celligner preserves and aligns biologically meaningful intra-cluster variability. For example, even though the melanoma tumors and cell lines mainly formed a single distinct cluster, variability within this cluster recapitulated recently-described melanoma differentiation states ^41^, and annotations of these melanoma subtypes were well-aligned between cell lines and tumors (**Fig. 4e**). Interestingly, the region of the melanoma cluster that consisted entirely of tumor samples primarily contained tumors of the transitory subtype that are from primary, rather than metastatic, samples. This result is consistent with the fact that many of the melanoma cell lines are annotated as being derived from metastatic samples ^1^. Together, these results highlight the ability of Celligner to reveal more detailed patterns of transcriptional similarity between cell lines and tumors, going beyond merely matching clusters.

One potential concern with methods that seek to globally align tumor and cell line data is that they might obscure important underlying biological differences. A key feature of Celligner in this regard is that it allows for sub-populations that are only present in one dataset or the other. For example, both data sets contain renal cancer samples, but samples annotated as chromophobe renal cell carcinoma are only present in the tumor data. Accordingly, after the Celligner alignment, the cluster of chromophobe renal cell carcinoma tumor samples remained distinct and did not include any cell lines (**Fig. 4f**). Similarly, myeloma cancer samples are present in the cell line data, but not in the TCGA, TARGET or Treehouse tumor data. After Celligner correction, the myeloma cell lines clustered near the other hematopoietic samples, but did not clearly group with any tumor samples (**Fig. 4d**). These cases demonstrate that Celligner does not artificially force all samples to align with samples from the other dataset, allowing it to reveal subtypes that are absent or underrepresented in either dataset.

### Information transfer between cell line and tumor datasets

By providing an unsupervised data integration procedure, Celligner enables joint analyses of the tumor and cell line datasets, providing greater power to detect transcriptionally distinct subpopulations. This is particularly true for the cell line data where there are ∼10-fold fewer samples compared to the tumors. Indeed, clustering analysis of the Celligner-aligned integrated dataset revealed a larger number of more distinct clusters among the cell lines compared with the same analysis applied to the cell line dataset on its own (**Supplementary Fig. 1**). This difference was most evident for cancer types that had few representative cell lines. For example, in the current dataset, only one testicular cell line is present (SUSA), and when analyzing the cell line data on its own, this cell line clustered most closely with the soft tissue cancers (**Supplementary Fig. 5**). Joint analysis of the Celligner-aligned data, however, showed that this cell line clustered with a subset of the germ cell tumors (**Supplementary Fig. 5**), and in particular was nearest neighbors with the non-seminoma testicular cancer samples (**Supplementary Fig. 5**).

We next explored whether integrated analysis of the tumor and cell line data might also help resolve missing, or potentially incorrect (sub)type annotations. For example, four cell lines that are not annotated as melanoma samples nevertheless clustered with the melanoma samples (**Supplementary Fig. 5**). One such cell line, COLO699, is annotated as derived from a metastatic lung cancer sample ^1^, raising the possibility that the current annotation accurately characterizes the biopsy site, but not the primary tissue. Previous reports in the literature have also identified that this cell line likely derives from a melanoma sample ^11^.

We can also use the combined dataset to perform ‘label transfer’ of annotations from one dataset to another. For example, ALL subtype annotations (T-cell and B-cell) were available for the ALL cell lines, but only for some of the ALL tumor samples. The ALL cell lines formed two distinct clusters, which perfectly matched the labeled B-cell and T-cell subtype. The ALL tumors also largely clustered together with the ALL cell lines, with all of the annotated (B-cell) tumor samples clustering with the B-cell cell lines. The rest of the (un-annotated) tumor samples could easily be classified as either B-cell or T-cell ALL (with some putative AML samples as well) based on their cluster membership (**Fig. 4c**), which aligned well with clustering of the tumor samples based on expression of B-cell ALL and T-cell ALL marker genes (**Supplementary Fig. 5**) ^44^. These results further highlight the advantage of performing an unsupervised global alignment that does not rely on existing annotations.

### Discovery of a novel group of transcriptionally and functionally distinct cell lines

Jointly analyzing the Celligner-aligned cell line and tumor data also revealed common structures across cell lines that were distinct from primary tumors. As described above (**Fig. 3**), cell line models of certain cancer types, such as thyroid and CNS, did not recapitulate the disease-specific transcriptional patterns exhibited by primary tumors of their respective cancer types. Closer inspection revealed that 252 of the cell lines that did not group with tumor samples of the same disease type formed a separate cluster (**Fig. 5a**). While approximately 20% of the cell lines belonged to this cluster, less than 2% of the tumor samples (primarily soft tissue and bone tumors) belonged to this cluster (**Supplementary Table 1**). The cell lines in the cluster spun a wide range of different lineages (notably, 82% of all CNS, 91% of all thyroid lines, and 41% of all liver lines; **Fig. 5b; Supplementary Table 1**).

**Figure 5.**
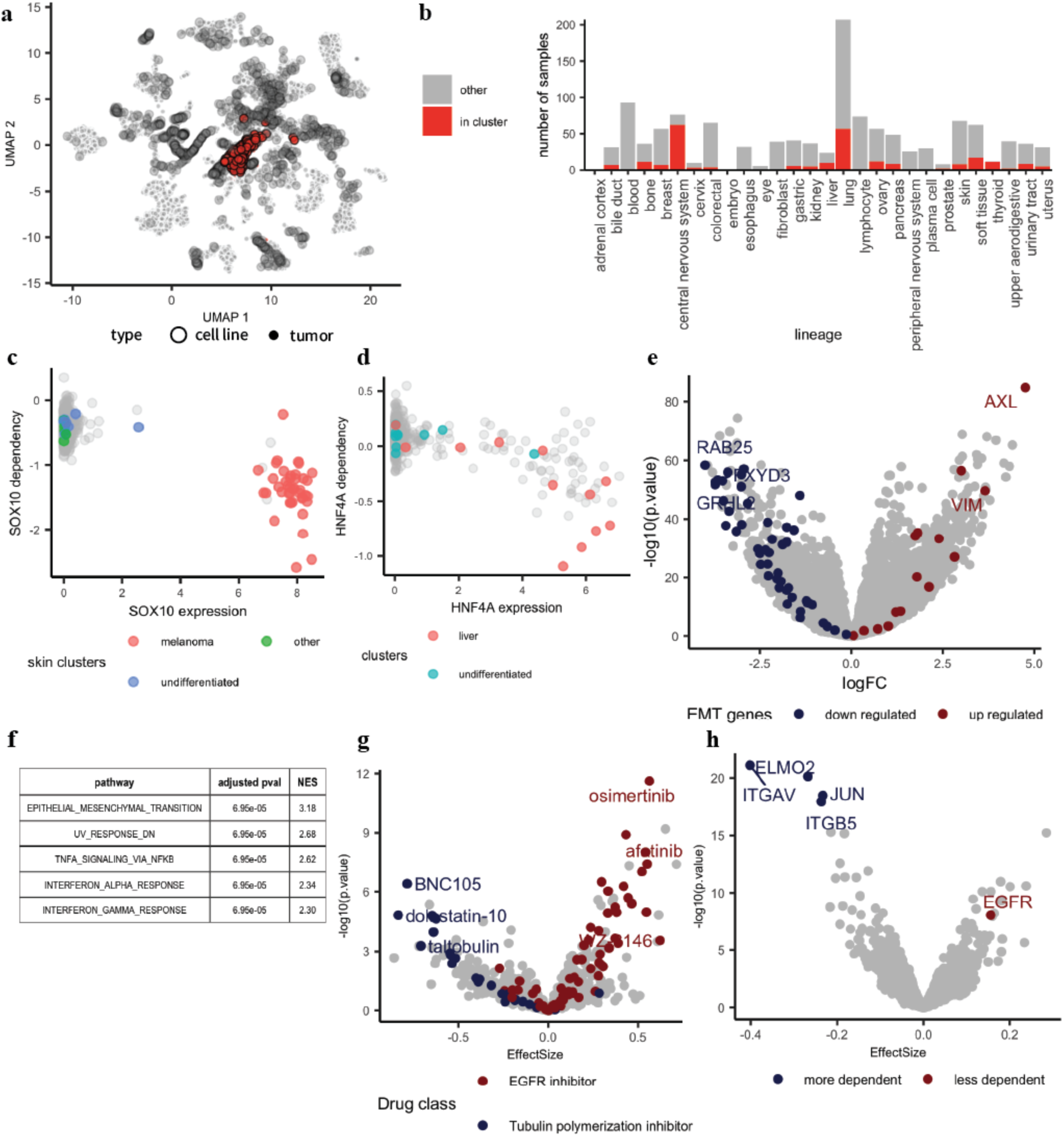
Cluster of cell lines show EMT signature and integrin-related dependencies. **a** A cluster of undifferentiated cell lines within the global Celligner-alignment. **b** Composition of the cell lines within the cluster. **c** Skin cell lines within the cluster are not dependent on SOX10. **d** Liver cell lines within the cluster are less dependent on HNF4A. **e** Differential expression analysis shows an up-regulated mesenchymal profile and **f** enrichment of the EMT pathway for cell lines in the cluster. **g** Differential drug vulnerability analysis shows decreased sensitivity to EGFR inhibitors and increased sensitivity to tubulin polymerization inhibitors for cell lines in the cluster. **h** Differential dependency analysis show stronger integrin related dependencies for cell lines within the cluster.

Cell lines in this cluster lacked lineage-specific expression characteristics present in the primary tumor datasets analyzed herein, suggesting that they have entered a more undifferentiated state. Indeed, of the twelve skin samples in this cluster that were annotated by Tsoi et al., all three of the skin cell lines and seven of the nine skin tumors were annotated as being of an ‘undifferentiated’ subtype ^41^. The majority (11/12) of the thyroid cell lines, which have been observed to be more dedifferentiated than thyroid tumors ^34,45,46^, also belonged to this cluster. To further assess how distinct these cell line models are from their lineage-matched counterparts that co-clustered with tumors, we also looked at a set of lineage-specific transcription factors. For example, SOX10 is selectively expressed in melanoma cells, and SOX10 knockout by CRISPR is lethal selectively in melanoma cell lines ^3^. Consistent with the interpretation of this cell-line-specific cluster as representing a more de-differentiated state, melanoma lines within the cluster showed much weaker expression of, and dependency on, SOX10 (**Fig. 5c**) ^41^. Similarly, liver cancer cell lines within the ‘undifferentiated’ cluster showed lower expression of, and less dependency on, the hepatocyte transcription factor HNF4A (**Fig. 5d**).

To further understand the biological features that distinguish this group of undifferentiated cell lines we performed genome-wide differential expression analysis, controlling for differences attributable to the annotated lineages. This analysis revealed a striking enrichment of epithelial-mesenchymal transformation (EMT)-related genes (**Fig. 5e, f**), reflecting a stronger mesenchymal expression pattern among these undifferentiated cell lines. The few tumor samples in this cluster were primarily from cancer types with mesenchymal cell lineages ^47^ and GSEA of genes differentially expressed by these samples showed elevated expression of the EMT pathway (normalized enrichment score = 3.33, adjusted p-value = 6.2e-05).

We also tested whether the cell lines expressing this distinct mesenchymal/undifferentiated expression pattern exhibit a unique pattern of chemical and genetic vulnerabilities. For this, we used the Achilles dataset of genome-wide CRISPR knockout screens to interrogate gene essentiality across 689 cell lines ^48^, as well as a recently-generated dataset of clinical compounds screened across 578 cell lines ^5^. Examination of differences in the drug sensitivities of these undifferentiated cell lines showed that they have increased sensitivity to tubulin polymerization inhibitors (**Fig. 5g**), as well as greater dependency on integrin genes, particularly ITGAV and ITGB5 (**Fig. 5h**), consistent with their upregulation of EMT-related genes ^49,50^. The undifferentiated cell lines were also more resistant than other cell lines to several compounds, most notably many EGFR inhibitors (**Fig. 5g**). This is also consistent with a marked decrease in EGFR dependency among the undifferentiated cell lines (**Fig. 5h**).

## Discussion

Cancer cell lines are crucial drivers of preclinical cancer research. Yet, our limited understanding of the similarities and differences between cell lines and patient tumors remains a key challenge for translating findings from cell lines to the clinic. To help address this, we developed a computational method, Celligner, which identifies and removes systematic differences between the RNA-Seq gene expression profiles of tumors and cell lines in an unsupervised manner, allowing for direct and detailed comparisons of the transcriptional states of cell lines and tumors.

In our global analysis, we identified pronounced differences across cancer types in how well cancer cell lines reflected the transcriptional patterns of their primary tumor counterparts. While many disease types (such as lymphoma, Ewing sarcoma, and melanoma) were similar between cell lines and tumors, there were few thyroid and CNS cell lines whose gene expression profiles aligned with the corresponding primary tumor samples. Previous studies have identified that CNS cell lines grown in serum-containing media tend to lose their ability to differentiate, and have gene expression profiles that are unlike their primary tumors ^40,51^, while CNS cell lines grown in serum-free specialized media had gene expression profiles and genetic aberrations that better recapitulated their primary tumors ^51^.

These differences pinpoint where new cell lines, organoid models ^52^, patient-derived xenografts and mouse models are most needed. They also reinforce the importance of efforts such as the Human Cancer Models Initiative (HCMI) that aim to address gaps in our current *in vitro* model representation. Future applications of the method to RNAseq datasets from these and other novel model formats should prove useful.

We discovered a distinct set of cell lines, composed of a range of tissue types, which exhibited a transcriptional state that was largely dissimilar from those of the available primary tumor samples. These cell lines had undifferentiated characteristics, lacking activity of lineage-specific transcription factors (and associated genetic dependencies). They also showed clear upregulation of mesenchymal genes and downregulation of epithelial markers; all characteristics concordant with an EMT phenotype.

Consistently, this group of cell lines was most similar to tumors that arise from mesenchymal tissue, but generally clustered separately from the primary tumor samples. Interestingly, these undifferentiated cell lines also exhibited distinct genetic and chemical vulnerabilities, including increased dependency on integrin genes and sensitivity to tubulin inhibitors, as well as decreased sensitivity to EGFR inhibitors, all of which are consistent with an EMT state ^53–55^. This raises the possibility that these cell lines may reflect a biologically relevant tumor cell state that is not represented in the primary tumor datasets used here. Indeed, the EMT program has been most clearly identified in early metastatic samples ^56^, while the tumor data we used is largely from primary tumor samples ^10^. More research is needed to determine whether these cell lines could be good models for tumors that have undergone EMT, or if they reflect an artifact of cell culture conditions. As new large-scale datasets of metastatic and drug-resistant tumors emerge we can incorporate them into Celligner to better answer this question.

A recent comparison of CCLE and TCGA expression data by Yu et al. used a combination of COMBAT correction and linear regression to remove expression patterns associated with tumor purity ^20^. They used their analysis to rank cell lines by tumor type and establish a panel of 110 cell lines across 22 cancer types that they identified as good representatives of tumors. Our unsupervised approach identified similar differences between cell lines and tumors, such as immune signatures and cell cycle differences. Our analysis was also able to identify substantial disease-specific differences in how well cell line models reflected tumor transcriptional states ^39,40^, as well as revealing a large group of (to our knowledge previously undescribed) ‘undifferentiated’ cell lines, as detailed above.

Our analyses focused on using gene expression data to compare tumor and cell line samples. In contrast, previous efforts, such as Cellector ^15^, have utilized genomic alterations to identify cell lines that are most representative of specific disease subtypes. In general, we found that cell lines previously identified as poor models based on copy number and mutations were often identified as non-tumor-like based on our analysis of Celligner-aligned gene expression features as well. For example, Domcke et al. observed that OC316 was hyper-mutated ^12^, Sinha et al. found that SLR20 had an outlier copy number profile ^57^, and Ronen et al. found that COLO320 was dissimilar to colorectal tumors and lacked major colorectal cancer driver genes ^58^. In our analysis, all of these cell lines were also identified as being unlike their respective tumor types. This may reflect the fact that genomic alterations also result in corresponding changes in gene expression ^59,60^. In general, the use of gene expression profiles for comparing cancer samples is advantageous as it provides an information-rich readout of cell state, does not require predefined sets of genomic lesions, and avoids the need for matched normal samples. However, a future version of Celligner that also integrates genomic features could enable more detailed comparisons of tumors and cell lines.

A key component of Celligner is correcting for systematic differences between tumor and cell line expression profiles, most notably those related to contaminating normal cells in tumor samples. To do this, we utilized an unsupervised approach that did not depend on pre-defined signatures of the various contaminating cell types, and that also allowed us to account for unknown systematic differences between tumors and cell lines. For instance, we found that cell lines exhibited upregulation of cell cycle expression programs compared with tumors (**Supplementary Fig. 3**), which agrees with previous findings that a higher proportion of cancer cells are cycling *in vitro* compared to *in vivo* ^*61*^. The tumor/cell line differences we identified (MNN correction vectors) also varied across disease type, emphasizing the importance of using a non-linear method that allows for disease-type-specific differences. As single-cell data from normal tissues become more readily available, methods that use these data to estimate and remove the effect of different contaminating cells ^62,63^ could be incorporated to further improve comparisons between tumors and cell lines.

In order to facilitate the use of Celligner, we have incorporated an interactive web app on the Cancer Dependency Map portal (https://depmap.org/portal/celligner), that allows users to explore a Celligner-aligned integrated resource of cell line and tumor expression profiles, as well as download the data. This tool enables the identification of cell line models that best represent the transcriptional features of a tumor type, or even a particular tumor sample, of interest. More generally, by identifying and removing many of the confounding differences between cell lines and tumors in an unbiased fashion, Celligner enables integrated analyses of cell line and tumor datasets that can be used to reveal novel patterns within, and relationships between, these data, helping to improve translation of insights derived from cell line models to the clinic.

## Acknowledgements

The results shown here are in whole or part based upon data generated by the TCGA Research Network: https://www.cancer.gov/tcga. This work was supported in part by NCI U01 CA176058 (WCH) and NCI Cancer Model Development Center [Task Order Number HHSN26100008 under Contract No. HHSN261201500003I] (JSB).

## Author Contributions

AW, AT and JMM conceived and designed the study. AW and JMM wrote the analysis code. AW performed the analysis with help from AJ. TS, FV, JSB helped with interpretation. AW wrote the manuscript. WCH, JSB, FV, AT and JMM reviewed and revised the manuscript. AT and JMM supervised the study. WCH, JSB and FV acquired funds for the study

## Competing Interests

W.C.H. is a consultant for ThermoFisher, Solasta, MPM Capital, iTeos, Frontier Medicines, and Paraxel and is a Scientific Founder and serves on the Scientific Advisory Board (SAB) for KSQ Therapeutics. F.V. receives research support from Novo Ventures. A.T. is a consultant for Tango Therapeutics. All authors were partially funded by the Cancer Dependency Map Consortium, but no consortium member was involved in or influenced this study.

## Methods

### Expression data

Gene expression data for 12,236 tumor samples were taken from Treehouse Public Expression Dataset v10 obtained from Xena browser (https://xenabrowser.net). The data set compiled samples from the UCSC Treehouse Childhood Cancer Initiative, the Therapeutically Applicable Research to Generate Effective Treatments (TARGET) program, and The Cancer Genome Atlas (TCGA). Cell line gene expression data for 1,249 samples was taken from the DepMap Public 19Q4 file: CCLE_expression_full.csv downloaded from the Cancer Dependency Map portal (https://depmap.org/)^48^. All gene expression data was processed using the STAR-RSEM pipeline and is TPM log_2_ transformed (with a pseudocount of 1 added). Gene expression data were subset to 19,188 protein-coding genes that were present in both the tumor or cell line data.

### Celligner method

To remove sources of variation that are unique to one of the data sets and align the cell line and tumor data we used a multi-step process. First, we used contrastive principal component analysis (cPCA) ^35^ to identify correlated variability that is enriched in the tumor data compared to the cell line data, or vice-versa. In order to avoid identifying signatures related to differences in the cancer type or subtype compositions of the datasets we first clustered the tumor and cell line data separately and subtracted the average expression of each cluster from all samples in the cluster to estimate the average intra-cluster covariance for tumors and cell lines. The data sets were clustered in 70-dimensional PCA space using a shared nearest neighbor (SNN) based clustering method implemented in the Seurat R package ^64^, with a resolution parameter of 5. We then regressed out the first four cPCs (components which are higher variance in the tumor data) from both the tumor and cell line data (**Supplementary Fig. 6**).

We then performed mutual nearest neighbors (MNN) correction ^37^ on the data sets, using the cell line data as the reference dataset. To identify mutual nearest neighbors between the two datasets we used a set of genes that showed high between-cluster variance in each data set. Specifically, we used limma ^65^ to estimate the across-cluster variation in each gene’s expression within each dataset, using the empirical-Bayes moderated F-statistics as a metric of between-cluster variability. We used the union of the top 1000 genes from each data set with the lowest F-statistics (**Supplementary Table 2)**. We modified the MNN algorithm from the R package scran ^66^ to use different *k* values (the numbers of nearest neighbors to consider) for each data set, which was necessary to account for the much larger set of tumor samples used compared with cell lines. Specifically, we used a *k* value of 5 to identify nearest neighbors in the cell line data and a *k* value of 50 to identify nearest neighbors in the tumor data. We verified that the output was robust to modest changes in these parameters (**Supplementary Fig. 6**) and stable even if a tissue type was removed from one of the datasets (**Supplementary Fig. 6**).

### Measuring tumor/cell line similarity

To evaluate the similarity of cell lines to tumor samples we performed PCA on the Celligner-aligned data, then took the Euclidean distance between each cell line and tumor in PCA space (using 70 components). Cell lines were classified as a tumor type by identifying the most frequently occurring tumor type within each cell line’s 25 nearest tumor neighbors.

### Differential expression analysis

Differential expression analysis was performed on gene-level read count data using the ‘limma-trend’ pipeline ^65,67^. We first subsetted the data to genes that had a counts-per-million value greater than one in 10 or more samples. The data were normalized per sample using the ‘TMM’ method from the edgeR package ^68^, and transformed to log2 counts-per-million using the edgeR function ‘cpm’. Linear model analyses, with empirical-Bayes moderated estimates of standard error, were then used to identify genes whose expression was most associated with covariates of interest, such as disease type, or membership in a particular cluster. When analyzing differential gene expression related to the ‘undifferentiated’ cell line cluster (**Fig. 5e**), we included disease type as a covariate in the model. The differential dependency analysis and differential drug analysis were also performed using the limma pipeline ^65,67^ with empirical Bayes moderated t-stats for p-values and disease type included as a covariate.

### Dependency data

We used estimates of gene dependency taken from the Achilles genome-wide CRISPR-Cas9 KO data ^3^, 19Q4 release ^1^. Specifically, we used gene effect estimates based on the CERES algorithm, taken from the file gene_effect_corrected.csv from the Cancer Dependency Map portal (https://depmap.org) ^48^.

### Drug sensitivity data

Cell line drug sensitivity data were taken from a dataset of repurposing drugs screened with PRISM ^5^. For the PRISM dataset replicate-collapsed, log fold change data at a 2.5 µM dose from the secondary screen were used. Specifically, we used the ‘secondary-screen-replicate-collapsed-logfold-change’ and ‘secondary-screen-replicate-treatment-info’ vailable on the Cancer Dependency Map portal (https://depmap.org) and figshare ^69^. Annotations of compound mechanism of action (MOA) were also taken from ‘repurposing related drug annotations’ from the CLUE data library (clue.io/data).

### Gene set enrichment analysis

For gene set enrichment analysis of gene expression profiles we used the fgsea R package ^69^, using 100,000 permutations of gene-level values (log fold change values in **Fig. 5d**, contrastive principal component loadings in **Fig. 1e, Supplementary Fig. 2**, and average MNN correction vectors in **Supplementary Fig. 3**), to calculate normalized enrichment scores for gene sets from the ‘Hallmark’ and ‘GO_biological_proccesses’ gene set collections from MSigDB v6.2 ^70^.

### 2D embedding

To compute 2D embeddings of gene expression profiles (*e.g*. **Fig. 2; Supplementary Table 1**) we used the UMAP method ^38^, as implemented in the Seurat v3 package ^64^. The UMAP embedding was computed on the first 70 principal components, using Euclidean distance, with an ‘n.neighbors’ parameter of 10, and a ‘min.dist’ parameter of 0.5.

### Code availability

The full source code implementing the method and generating figures is made available at https://github.com/broadinstitute/Celligner_ms.

### Data availability

All datasets used to generate the results presented here are publicly available. The results of Celligner applied to the TCGA, TARGET, Treehouse, and CCLE datasets are available in a Fighare dataset at: https://figshare.com/articles/Celligner_data/11965269.

## Notes

https://depmap.org/portal/celligner

https://figshare.com/articles/Celligner_data/11965269

https://github.com/broadinstitute/Celligner_ms

