## Supplemental figures for "Global computational alignment of tumor and cell line transcriptional profiles"

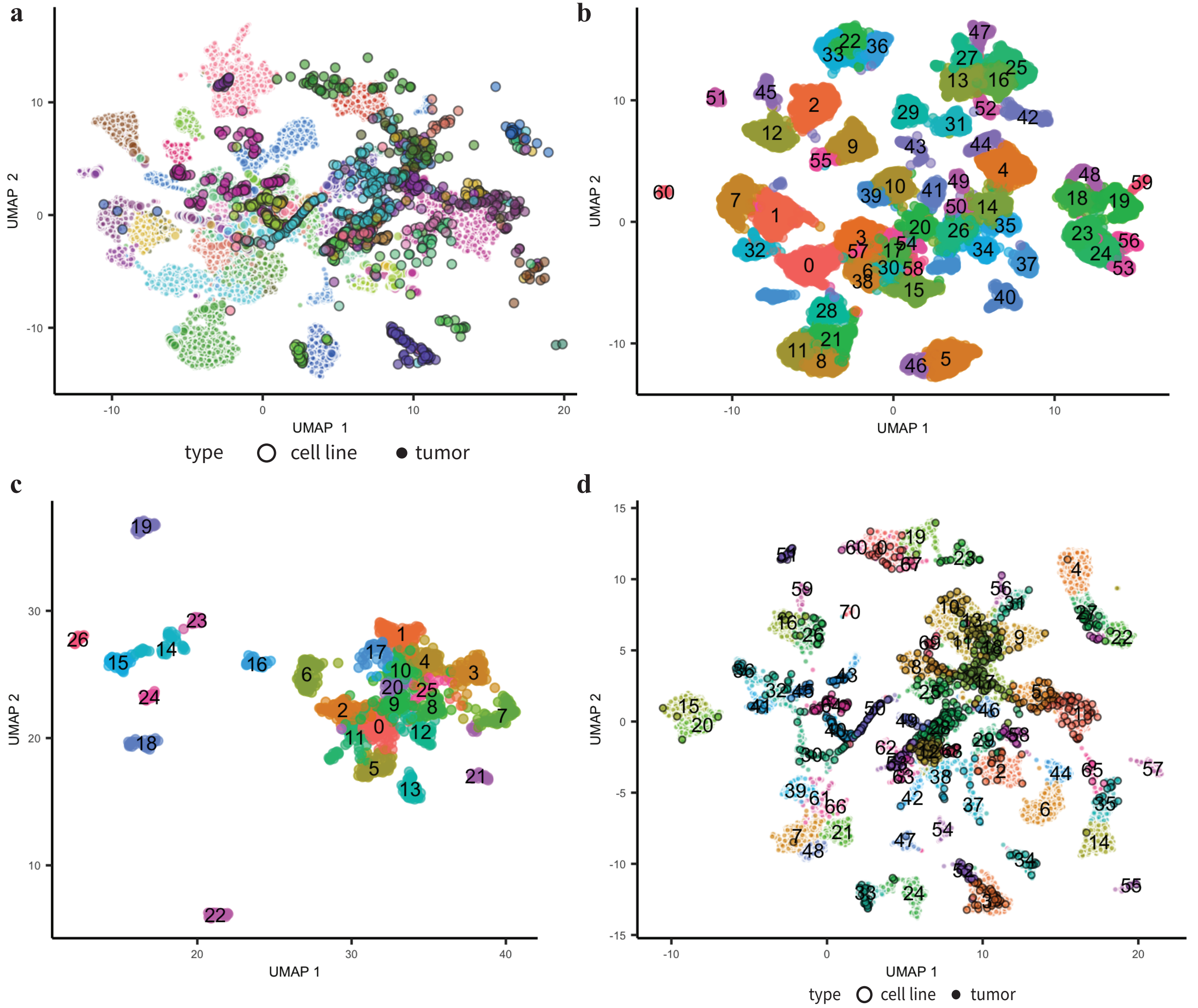


**Supplementary Figure 1. 2D Projections. a** Projection of cell lines and tumors after COMBAT correction colored by cancer lineage. Cancers of the same type do not align between cell lines and tumors. **b** Clustering of the uncorrected tumor expression data colored and labeled by the clusters identified. **c** Clustering of the uncorrected cell line expression data colored and labeled by the clusters identified. **d** Clustering of the Celligner-aligned tumor and cell line expression data colored and labeled by the clusters identified.

**a**

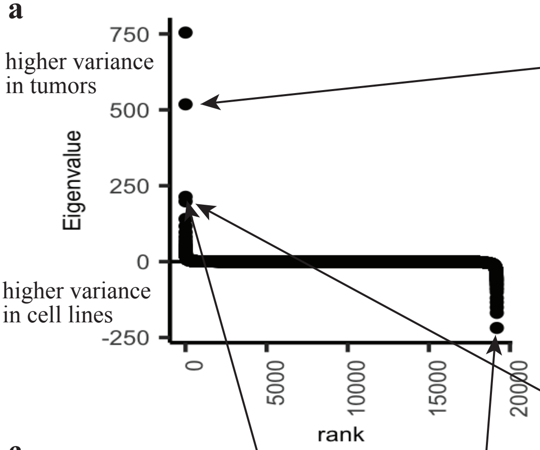


**b**

| pathway | adjusted pval | NES |
| --- | --- | --- |
| GO_ADAPTIVE_IMMUNE_RESPONSE | 0.000469 | 3.26 |
| GO_HUMORAL_IMMUNE_RESPONSE | 0.000469 | 3.15 |
| GO_LEUKOCYTE_MEDIATED_IMMUNITY | 0.000469 | 3.09 |
| GO_ADAPTIVE_IMMUNE_RESPONSE_BASED | 0.000469 | 3.07 |
| GO_LYMPHOCYTE_MEDIATED_IMMUNITY | 0.000469 | 3.07 |
| GO_RNA_SPLICING_VIA_TRANSESTERIFICATION | 0.000636 | -2.30 |
| GO_SISTER_CHROMATID_SEGREGATION | 0.000587 | -2.30 |
| GO_PEPTIDYL_LYSINE_MODIFICATION | 0.000647 | -2.26 |
| GO_RIBONUCLEOPROTEIN_COMPLEX_LOCALIZATION | 0.000556 | -2.25 |
| GO_RNA_LOCALIZATION | 0.000592 | -2.24 |

**c**

| pathway | adjusted pval | NES |
| --- | --- | --- |
| GO_ESTABLISHMENT_OF_PROTEIN_LOCALIZATION | 0.00129 | 1.93 |
| GO_MITOCHONDRIAL_RESPIRATORY_CHAIN_COMPLEX | 0.00129 | 1.90 |
| GO_PROTEIN_LOCALIZATION_TO_ENDOPLASMIC | 0.00129 | 1.88 |
| GO_NUCLEAR_TRANSCRIBED_MRNA_CATABOLIC | 0.00129 | 1.88 |
| GO_OXIDATIVE_PHOSPHORYLATION | 0.00129 | 1.85 |
| GO_MAGNESIUM_ION_TRANSMEMBRANE_TRANSPORT | 0.02460 | -2.25 |
| GO_ESTABLISHMENT_OF_PROTEIN_LOCALIZATION | 0.02310 | -2.19 |
| GO_MAGNESIUM_ION_TRANSPORT | 0.05230 | -2.18 |
| GO_PHAGOCYTOSIS_RECOGNITION | 0.02780 | -2.10 |
| GO_CHROMATIN_SILENCING_AT_RDNA | 0.04250 | -1.97 |

**d**

| pathway | adjusted pval | NES |
| --- | --- | --- |
| GO_EXTRACELLULAR_STRUCTURE_ORGANIZATION | 0.000177 | 2.87 |
| GO_MUSCLE_CONTRACTION | 0.000177 | 2.76 |
| GO_MUSCLE_SYSTEM_PROCESS | 0.000177 | 2.72 |
| GO_REGULATION_OF_OSSIFICATION | 0.000177 | 2.67 |
| GO_MUSCLE_ORGAN_DEVELOPMENT | 0.000177 | 2.64 |
| GO_SISTER_CHROMATID_SEGREGATION | 0.000177 | -3.01 |
| GO_NUCLEAR_CHROMOSOME_SEGREGATION | 0.000177 | -2.94 |
| GO_CHROMOSOME_SEGREGATION | 0.000177 | -2.94 |
| GO_SISTER_CHROMATID_COHESION | 0.000177 | -2.83 |
| GO_MITOTIC_SISTER_CHROMATID_SEGREGATION | 0.000177 | -2.73 |

**e**

| pathway | adjusted pval | NES |
| --- | --- | --- |
| GO_RESPONSE_TO_TYPE_I | 0.000541 | 2.80 |
| GO_DEFENSE_RESPONSE_TO_VIRUS | 0.000541 | 2.50 |
| GO_NEGATIVE_REGULATION_OF_VIRAL | 0.000541 | 2.45 |
| GO_NEGATIVE_REGULATION_OF_VIRAL | 0.000541 | 2.43 |
| GO_INTERFERON_GAMMA_MEDIATED_SIGNALING | 0.000541 | 2.42 |
| GO_DNA_DEPENDENT_DNA_REPLICATION | 0.000924 | -2.59 |
| GO_RNA_SPLICING_VIA_TRANSESTERIFICATION | 0.001150 | -2.53 |
| GO_DNA_REPLICATION | 0.001040 | -2.52 |
| GO_DNA_RECOMBINATION | 0.001040 | -2.49 |
| GO_RIBONUCLEOPROTEIN_COMPLEX_LOCALIZATION | 0.000934 | -2.47 |

**Supplementary Figure 2. contrastive Principle Component Analysis (cPCA) a** cPCA eigenvalue spectrum. **b** GSEA of the second, **c** third, **d** and fourth cPCs, which are higher variance in tumors. **e** GSEA of the top cell line specific cPC.

a

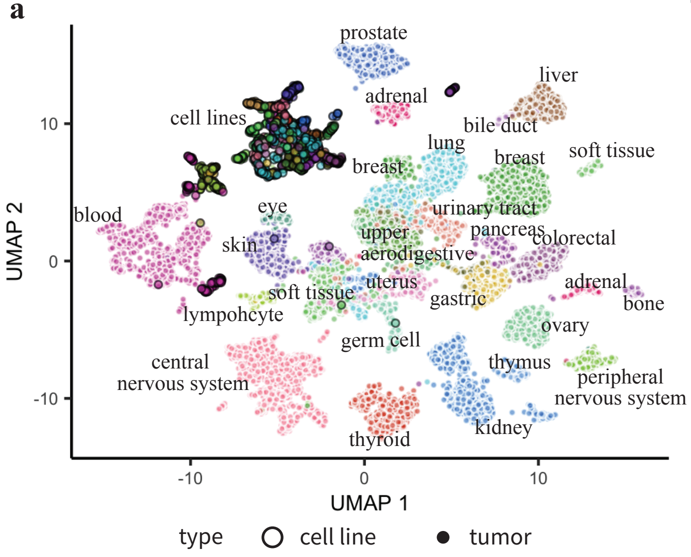


b

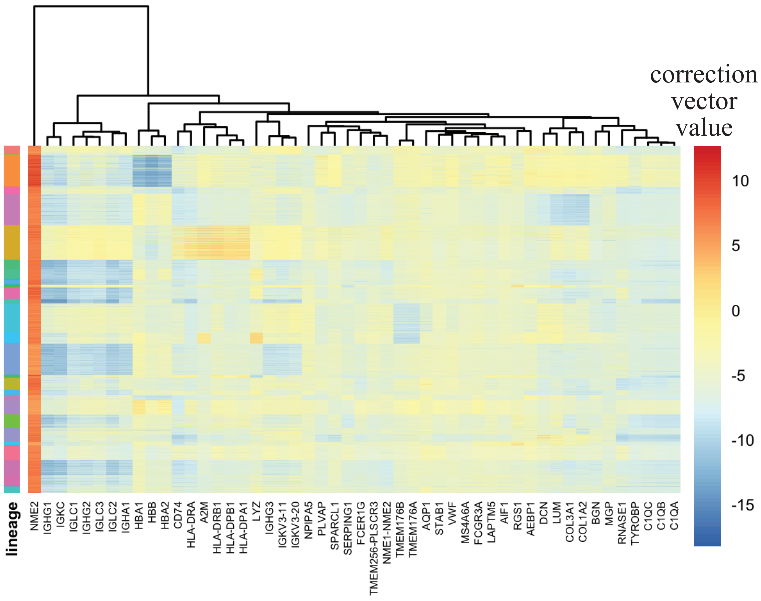


c

GSEA of average MNN correction vectors

| pathway | adjusted pval | NES | higher variance in |
|---|---|---|---|
| GO_HUMORAL_IMMUNE_RESPONSE | 0.000408 | -2.83 | tumors |
| GO_PROTEIN_ACTIVATION_CASCADE | 0.000408 | -2.78 | tumors |
| GO_COMPLEMENT_ACTIVATION | 0.000408 | -2.75 | tumors |
| GO_HUMORAL_IMMUNE_RESPONSE_MEDIATED | 0.000408 | -2.74 | tumors |
| GO_B_CELL_MEDIATED_IMMUNITY | 0.000408 | -2.70 | tumors |
| GO_SISTER_CHROMATID_SEGREGATION | 0.000574 | 3.12 | cell lines |
| GO_CHROMOSOME_SEGREGATION | 0.000635 | 3.08 | cell lines |
| GO_NUCLEAR_CHROMOSOME_SEGREGATION | 0.000597 | 3.08 | cell lines |
| GO_SISTER_CHROMATID_COHESION | 0.000547 | 2.91 | cell lines |
| GO_MITOTIC_SISTER_CHROMATID_SEGREGATION | 0.000537 | 2.90 | cell lines |

d

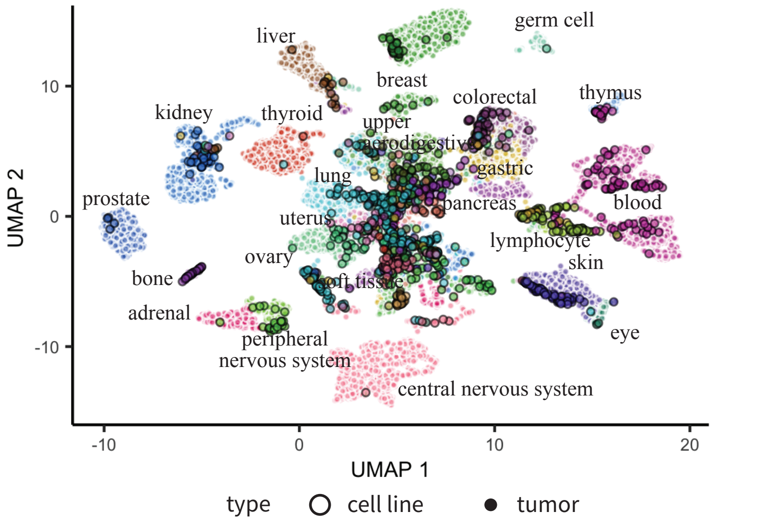


e

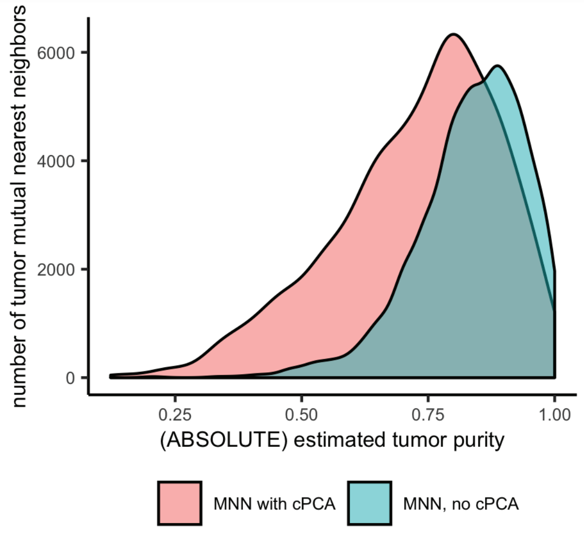


**Supplementary Figure 3. cPCA and MNN correction.** **a** Projection of cell lines and tumor using just cPCA correction colored by cancer lineage. **b** Heatmap of the MNN correction vectors, showing the 50 genes with the highest absolute average correction vector values. **c** GSEA of the average MNN correction vectors. **d** Projection of cell line and tumor data using just MNN correction colored by cancer type. **e** Running cPCA before MNN increases the number of identified mutual nearest neighbors, especially for lower purity tumors.

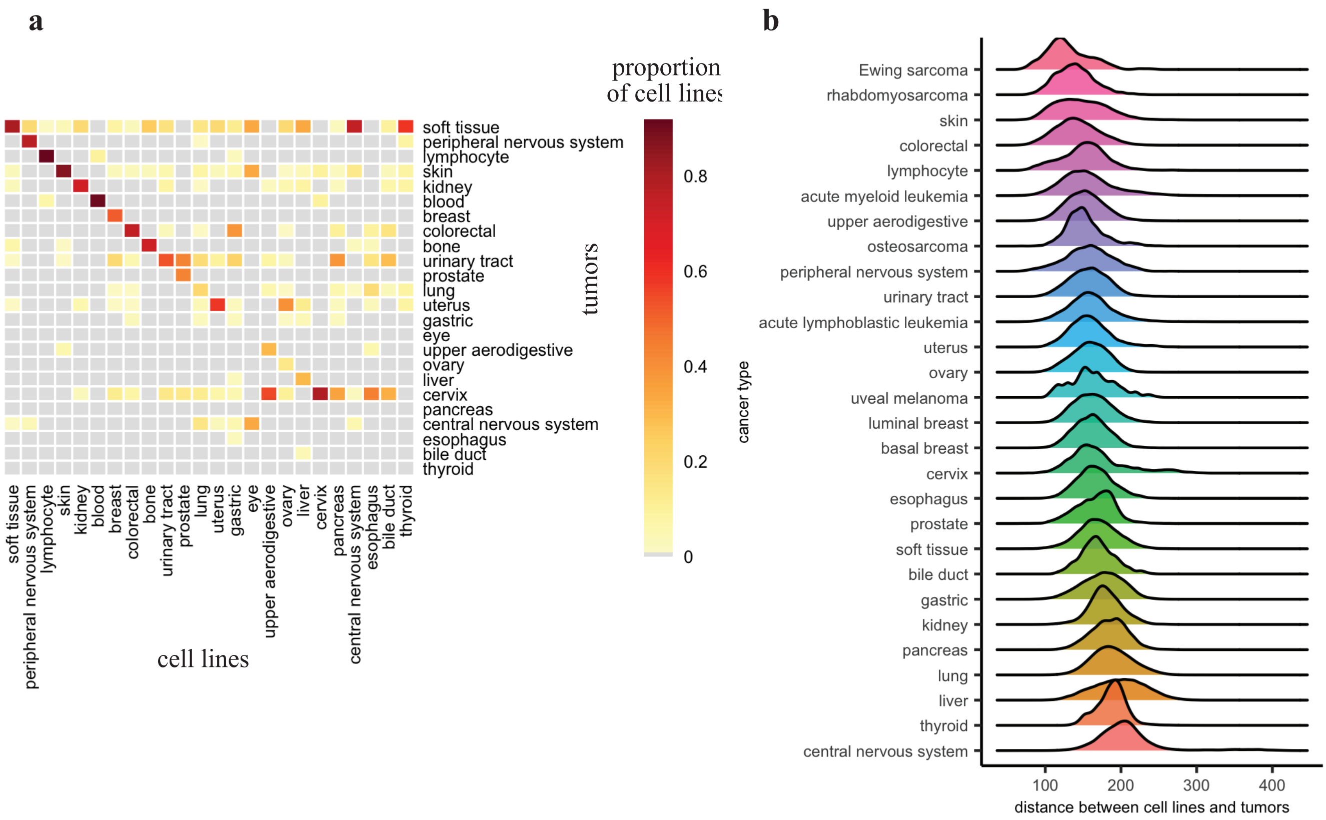


**Supplementary Figure 4. Uncorrected classification of cell lines by tumor type a** Proportion of cell lines that are classified as each tumor type using uncorrected data. 44% of cell lines with corresponding types in the tumor data matched to tumors of the same type. **b** Distribution of distances (using top 70 PCs) between cell lines and tumors of the same (sub)type using uncorrected data.

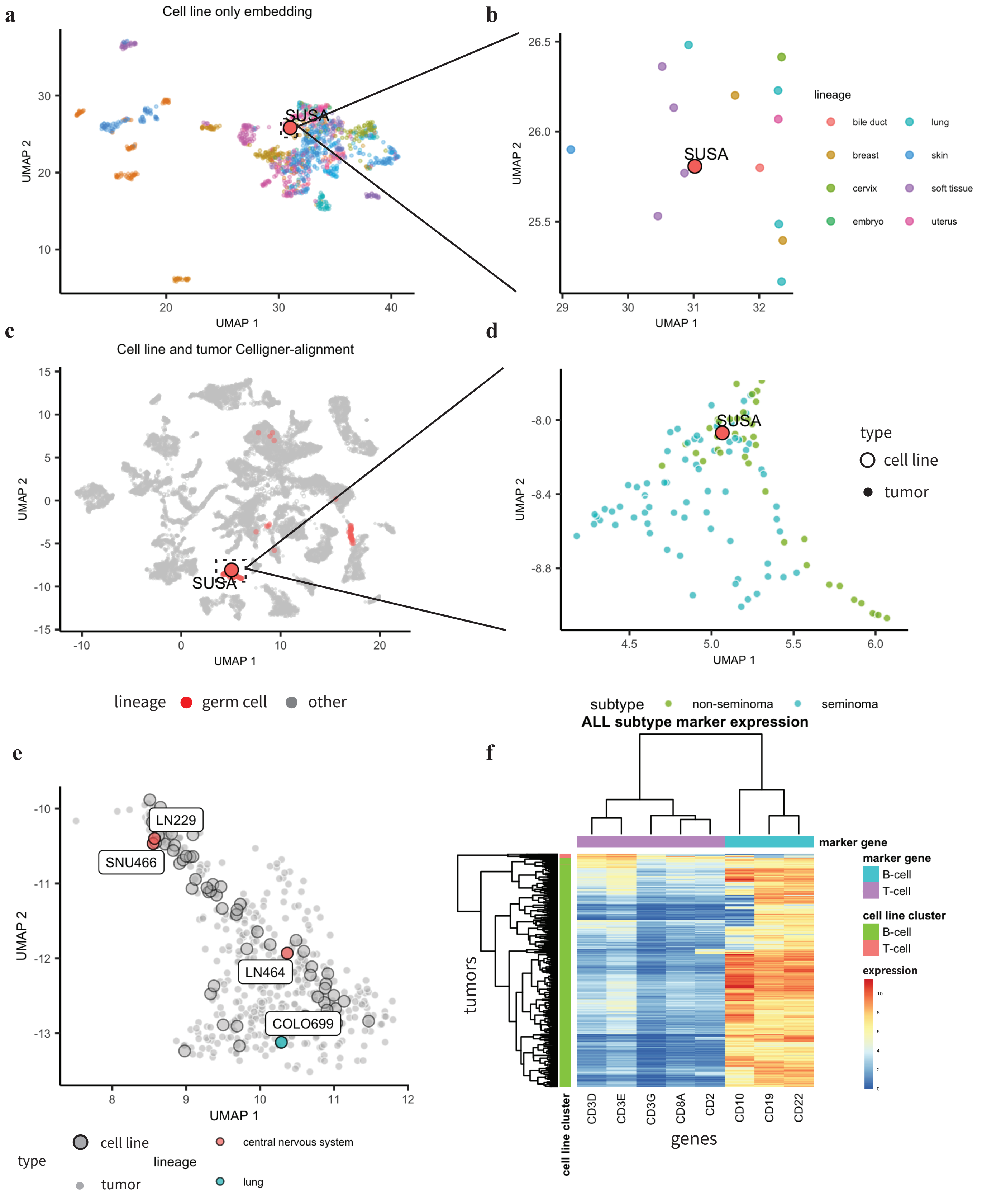


**Supplementary Figure 5. Integrated analysis of tumors and cell lines increases power to detect subtypes and identify misannotated samples.** We computed a 2D UMAP embedding of just the cell line RNA-Seq data and the Celligner-aligned data. **a, b** In the cell line only embedding SUSA, the only testicular cell line in the data, clusters near cell lines of a variety of tissue types, **c** while in the Celligner-aligned embedding SUSA clusters with the germ cell tumors, **d** and its nearest tumors neighbors are primarily non-seminoma testicular cancer samples. **e** Cell lines which are not annotated as skin cell lines, but cluster with the melanoma cluster. **f** Clustering of ALL tumor samples by expression of ALL subtype marker genes agrees well with the clustering of those samples with ALL T-cell and B-cell cell lines.

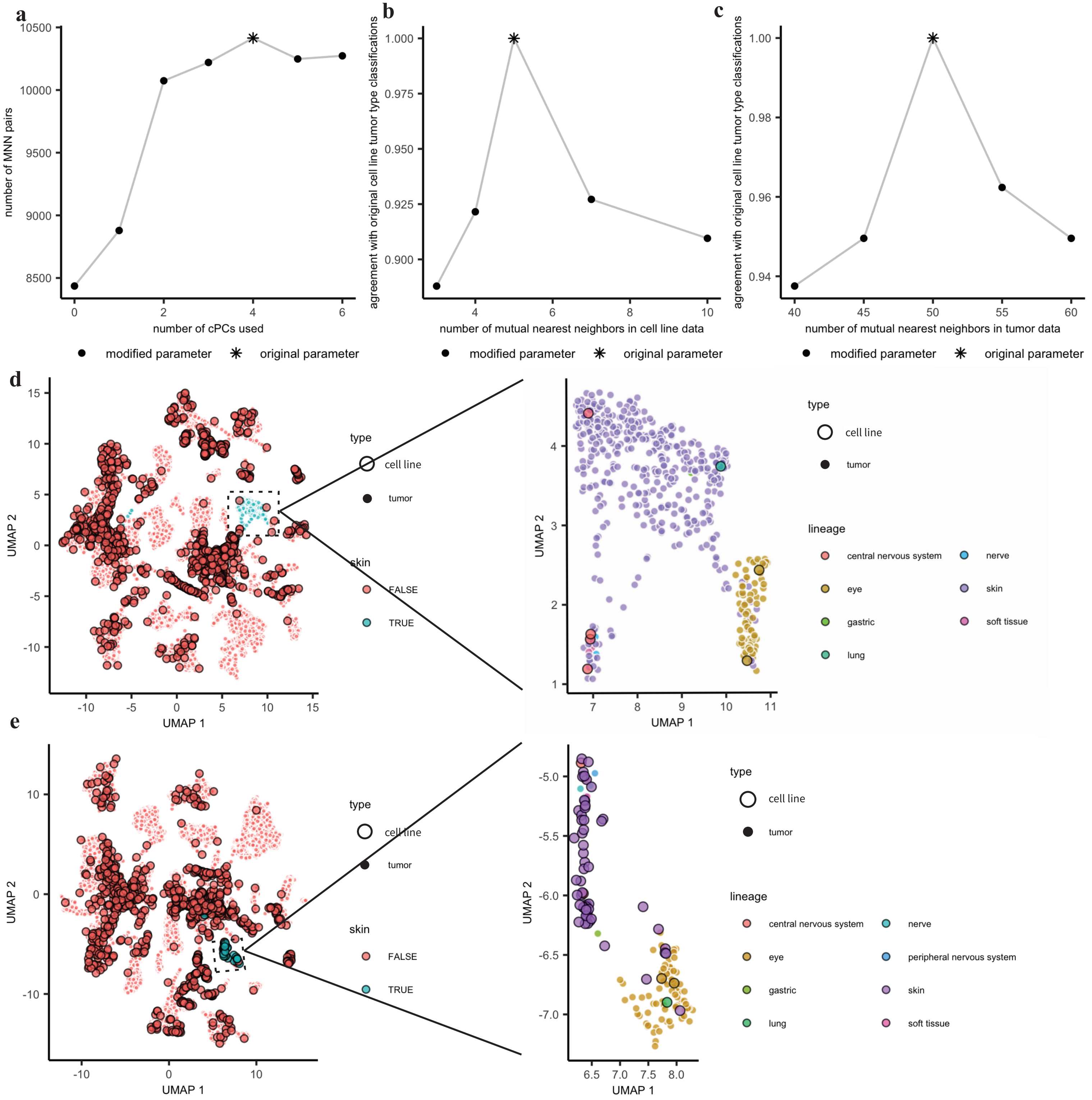


**Supplementary Figure 6. Celligner parameter selection and robustness. a** Removing contrastive principal components increases the number of mutual nearest neighbors. Regressing out the top four cPCs with higher variance in the tumor data increased the number of mutual nearest neighbors Celligner output is robust to alterations in parameters. **b, c** In order to evaluate the robustness of the output we varied one parameter at a time, keeping all other parameters fixed, then compared the tumor type classifications for each cell lines (see Methods) to those of the original. The parameters that we tested were the k parameters used as input to MNN correction. Celligner is robust to removal of data**.** We also tested the stability of the output to removal of a subset of the data. **d** We removed all of the cell lines annotated as skin and re-ran the alignment. We saw that even without these cell lines the skin tumors formed a clear cluster, and cell lines that we believe are mis-annotated continued to cluster with the skin tumors. **e** We also tested removing the skin tumors and re-running the alignment. In this case, the skin cell lines clustered near the uveal melanoma, but primarily formed a separate cluster without tumors.
